# Classification of the plant-associated phenotype of *Pseudomonas* strains using genome properties and machine learning

**DOI:** 10.1101/2021.07.30.454435

**Authors:** Wasin Poncheewin, Anne D. van Diepeningen, Theo AJ van der Lee, Maria Suarez-Diez, Peter J. Schaap

## Abstract

The rhizosphere, the region of soil surrounding roots of plants, is colonized by a unique population of Plant Growth Promoting Rhizobacteria (PGPR). By enhancing nutrient uptake from the soil and through modulation of plant phytohormone status and metabolism, PGPR can increase the stress tolerance, growth and yield of crop plants. Many important PGPR as well as plant pathogens belong to the genus *Pseudomonas*. There is, however, uncertainty on the divide between phytobeneficial and phytopathogenic strains as previously thought to be signifying genomic features have limited power to separate these strains. Here the Genome properties (GP) common biological pathways annotation system was applied to establish the relationship between the genome wide GP composition and the plant-associated phenotype of 91 *Pseudomonas* strains representing both phenotypes. GP enrichment analysis, Random Forest model fitting and feature selection revealed 28 discriminating features. A validation dataset of 67 new strains confirmed the importance of the selected features for classification. A number of unexpected discriminating features were found, suggesting involvement of novel molecular mechanisms. The results suggest that GP annotations provide a promising computational tool to better classify the plant-associated phenotype.

**Author summary:** With a growing population the need to double the agricultural food production is specified. Simultaneously, there is an urgent need to implement sustainable and climate change resilient agricultural practices that preserve natural ecosystems. Cooperative microbiomes play important positive roles in plant growth development and fitness. Properly tuned, these microbiomes can significantly reduce the need for synthetic fertilizers and can replace chemicals in crop pest control. To select beneficial candidates, their traits need to be described and likewise, potential detrimental traits should be avoided. Here we applied GP-based comparative functional genomics, enrichment analysis and Random Forest model fitting to compare known phytobeneficial and phytopathogenic *Pseudomonas* strains. A number of unexpected discriminating features were found suggesting the involvement of novel molecular mechanisms.

## Introduction

Among the targets set by the UN to achieve the zero-hunger goal, the need to double the agricultural food production is specified [1]. Earlier attempts to improve plant performance and production focused on plant breeding, pest control by chemical means and the implementation of synthetic fertilizers tapping into finite global reserves. While these strategies were successful in enhancing production, the increasing adverse effects on the environment challenges us to find sustainable alternatives [2–4].

A multitude of studies has demonstrated that cooperative microbiomes can play important positive roles in plant growth, development and fitness. One particular hotspot is the rhizosphere, the region of soil surrounding plant roots, colonized by Plant Growth Promoting Rhizobacteria (PGPR)[5]. A stable PGPR population can increase the stress tolerance, growth and yield of crop plants by enhancing nutrient uptake from the soil and through modulation of plant phytohormone status and metabolism [6–13]. As a result, a large catalogue of plant beneficial bacterial strains has been identified. The most studied are *Pseudomonas* spp., a functionally diverse group representing both plant beneficial and pathogenic strains [14–16].

A diverse spectrum of plant-host interaction pathways determines the plant-associated phenotype of a *Pseudomonas* strain. Correlational approaches have identified a number of marker genes contributing to the phenotype [17–19]. These genes are however, to a certain degree, shared between beneficial and pathogenic strains [20] and consequently, with each new genome addition the uncertainty on the divide between beneficial and the pathogenic strains increases. Until now, a generic description of presence and completeness of biological pathways contributing to the plant-associated phenotype of a *Pseudomonas* strain is lacking. Such knowledge would bring fundamental insights into their potential to enhance plant performance and resilience. When genes are placed in context of biological pathways comparative functional genomics is possible. Genome Properties (GP) is an annotation system whereby functional attributes can be assigned to a genome [21]. The resource represents a collection of 1286 common biological pathways, and each GP is evidenced by a distinct set of protein domains.

Here we applied GP-based comparative functional genomics to compare known phytobeneficial and phytopathogenic *Pseudomonas* strains using both traditional statistical analysis and machine learning methods. This allowed us to accurately classify *Pseudomonas* strains, and to identify discriminating features for both the phytobeneficial and phytopathogenic lifestyle. In the discussion section these discriminating features are placed into biological context.

## Results

Based on literature review, the complete genomes of 84 *Pseudomonas* strains were retrieved from the Pseudomonas Genome DB (version 17.2) [22] and categorized as encoding either a ‘phytobeneficial’ strain (51 strains) or a ‘phytopathogenic’ strain (33 strains). This selection was supplemented with the complete genomes of seven new or re-sequenced phytobeneficial strains; *P. putida* P9, *P. corrugata* IDV1, *P. fluorescens* R1 and WCS374, *P. protegens* Pf-5, *P. chlororaphis* Phz24 and *P. jessenii* RU47. To avoid gene and protein domain annotation inequality, the genome sequences of all 91 strains were *de novo* annotated. Subsequently, the two groups were compared using nucleotide sequence similarity, by protein domain presence and by presence and completeness of domain-based GPs (**Fig 1**). Domain content was subjected to enrichment analysis and the GP content of both groups was used to train a Random Forest (RF) model for classification and feature selection [23]. The performance of the classification methods was further validated using a set of 67 newly sequenced soil derived *Pseudomonas* genomes obtained from a newer version (V20.2) of the Pseudomonas Genome DB. Based on literature data. Using literature data, 17 strains of this validation set could be classified as phytobeneficial strains while 34 strains were involved in bioremediation. For 16 strains the classification was unclear however, a number of these strains were *P. chlororaphis* strains known to be phytobeneficial.

**Fig 1:**
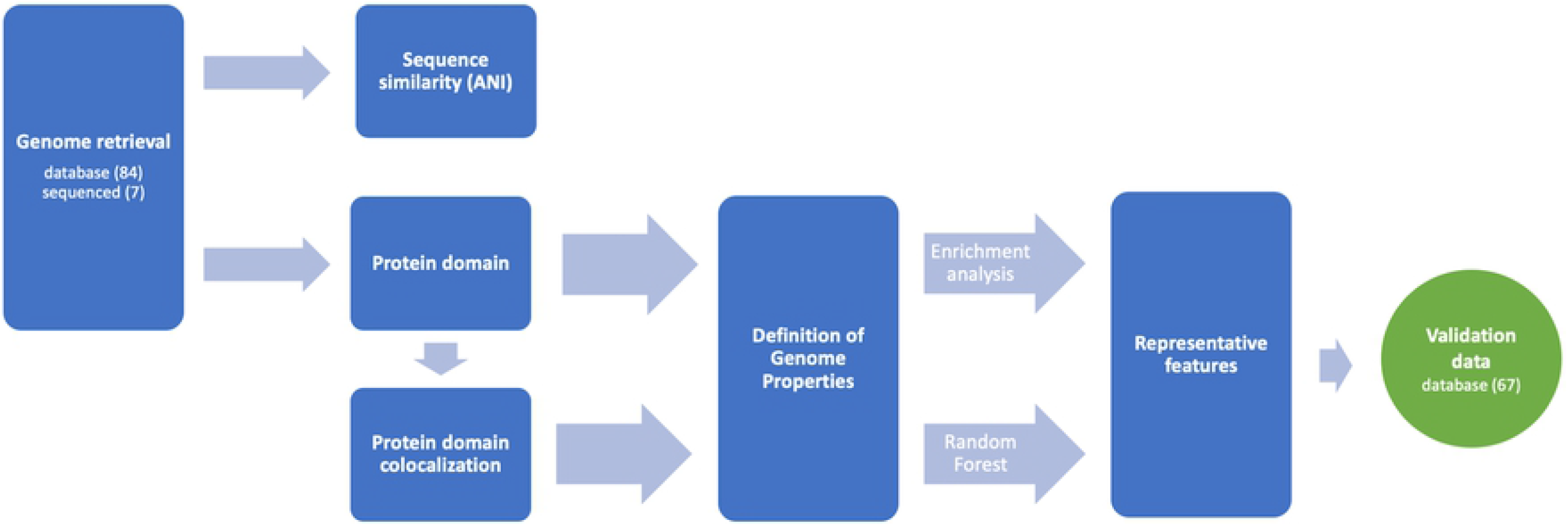
Workflow for GPs based functional genomics and classification. Genome sequences are analyzed using sequence similarity and protein domain content. (Colocalized) protein domain content is used to infer Genome Properties. Enrichment analysis and Random Forest feature selection was used obtain genomic features. Classification performance was evaluated using a validation dataset of 67 newly available genomes.

### Sequence similarity

We first examined the genomic relatedness between the phytobeneficial and phytopathogenic group, by calculating the Average Nucleotide Identity (ANI) scores between all possible pairs (**Fig 2**). The ANI scores showed that corresponding with their phenotypic classification the genome sequences could be divided into two groups with *Pseudomonas sp*. M30-35 being less similar to the rest of the phytobeneficial group. The average sequence similarity within the phytopathogen and the phytobeneficial group was 90.01 ± 5.53 and 79.57 ± 4.27 respectively. The ANI-score measures genomic similarity between the coding regions of two genomes at nucleotide-level taking into account hits that have 70% or more identity and at least 70% coverage of the shorter gene. The ANI score does not take into account the fraction of coding sequences that actually contribute to this score and thus provides no insight in the degree of strain-specific functional adaptations. To study which strain-specific functional adaptations impact the phenotype, the protein domain content of each strain was considered.

**Fig 2:**
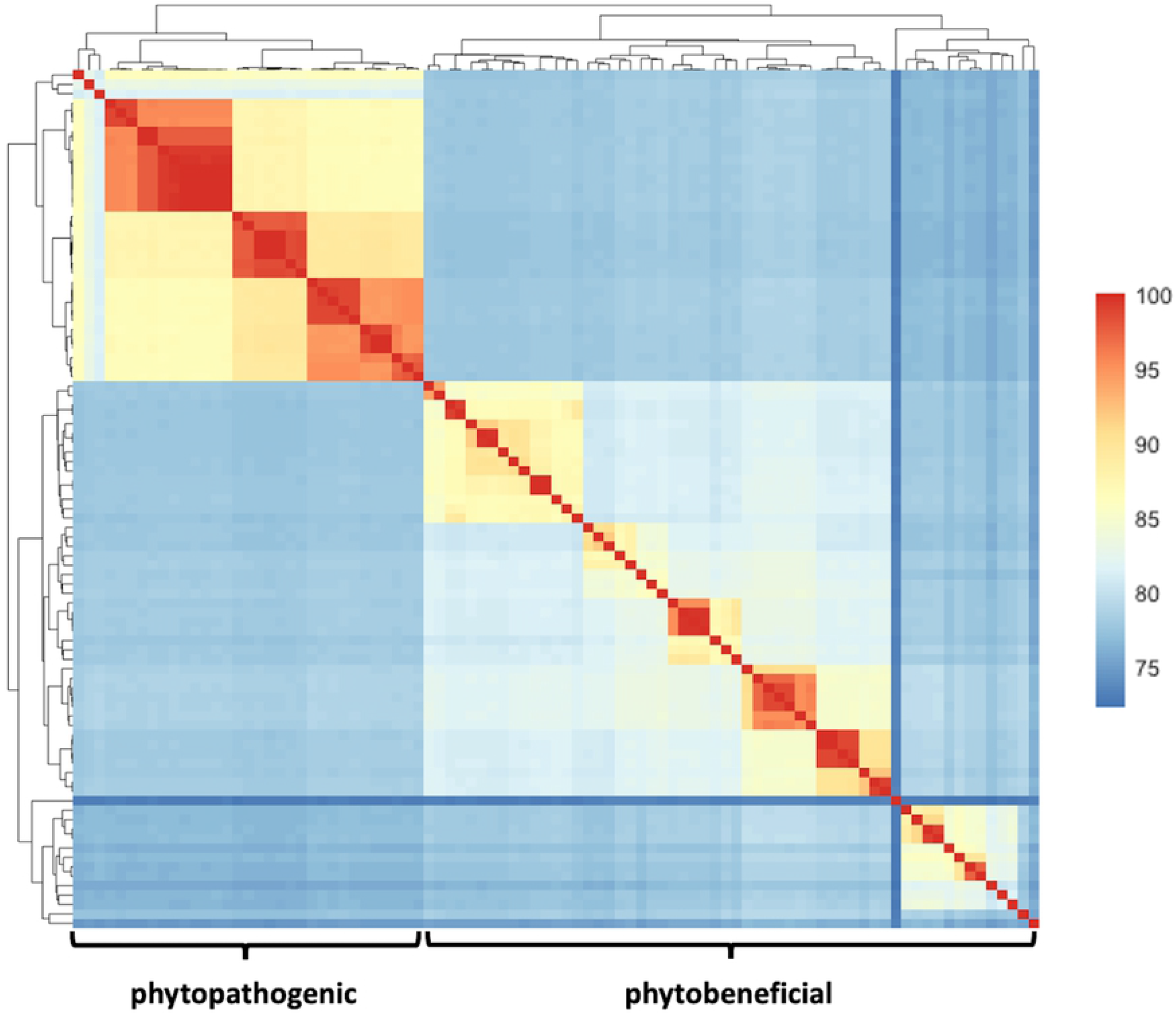
Pairwise Average Nucleotide Identity (ANI) scores between coding regions. Scores were calculated from alignments that have 70% or more identity and at least 70% coverage of the shorter gene.

### Protein domain content

The 91 complete Pseudomonas genomes contained, on average, 5640 ± 643 protein encoding genes. For each genome, 9342 ± 709 domains were identified with an average domain copy number of 2.35 ± 0.12 (**S1 Table**). Using domain presence as input, a group-wise enrichment analysis was done and a total of 410 and 329 protein domains were found to be significantly enriched in respectively phytobeneficial and phytopathogenic strains (**S2 Table**).

Phytobeneficial strains were enriched with five domains linked to Type II secretion systems (T2SS), ten domains linked to the term “cytochrome”, eight domains linked to, “quinohemoprotein” and six domains linked to “biofilm” (Poly-beta-1,6-N-acetyl-D-glucosamine type) biosynthesis. Interestingly, domains related to “quinohemoprotein” and “biofilm” were not only enriched but also exclusively found in phytobeneficial strains.

Phytopathogenic strains were enriched with domains involved in various types of secretion systems. Moreover, some of these domains were not present in any of the phytobeneficial strains. Eighteen of those pathogen enriched domains are reported to be involved in the Type III secretion system and five in the Type IV secretion system. In addition, the phytopathogen list showed enrichment of nine different domain involved in phosphonate metabolism. Functional clustering of enriched domains was further explored using genome properties.

### Genome properties

Genome properties (GP) represent a collection of currently 1286 common biological pathways. Each GP is constructed from a precomputed cluster of core protein domains which are used as essential evidence for the presence of the biological pathway [21]. Genome derived protein domains were used to construct for each strain a list of GPs with two possible evidence values: ‘YES’ indicating that the complete set of precomputed evidences had been detected and ‘PARTIAL’, indicating that the GP is likely present due to the presence of an incomplete set of evidences above a per GP specified minimal threshold. In addition, we took into account that the bacterial genes encoding domains that function in the same biological pathway are often arranged in operonic structures corresponding to syntenic blocks. For each strain therefore GPs were reconstructed not only based on protein domain presence/absence (GP-PA) but also on protein domain colocalization (GP-SND; synteny-non-directional) and on domain colocalization and being encoded on the same strand (GP-SD; synteny-directional). For domain colocalization a nearest neighbor approach was applied using a sliding window of 20 protein domains.

**Table 1** summarizes the results obtained for the three approaches.

A total of 438 GPs were not present in any the investigated Pseudomonas strains. The majority of these GPs represented functions and processes typically found in eukaryotic species (**S3 Table**). Conversely, using the GP-PA method, a functional GP core of 154 complete GPs present in all strains could be obtained. When domain colocalization was used as an additional constraint a functional core of 37 complete, likely operonic, GPs was found with both domain colocalization methods. Note that overall, the GP-SND and GP-SD generated very similar output underpinning a strong linkage between operonic structures and functional genome properties in bacterial species (**Table 1**).

**Table 1:**
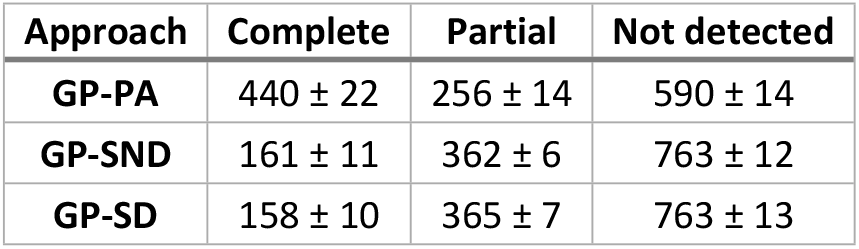
Number of strain specific GP classes per approach.

Next, a principal component analysis (PCA) was applied to the GP data. With all three methods a clear separation between the pathogen and the biocontrol group were obtained (**S4 Fig**). **Fig 3** shows the results obtained with the GP-SND approach.

**Fig 3:**
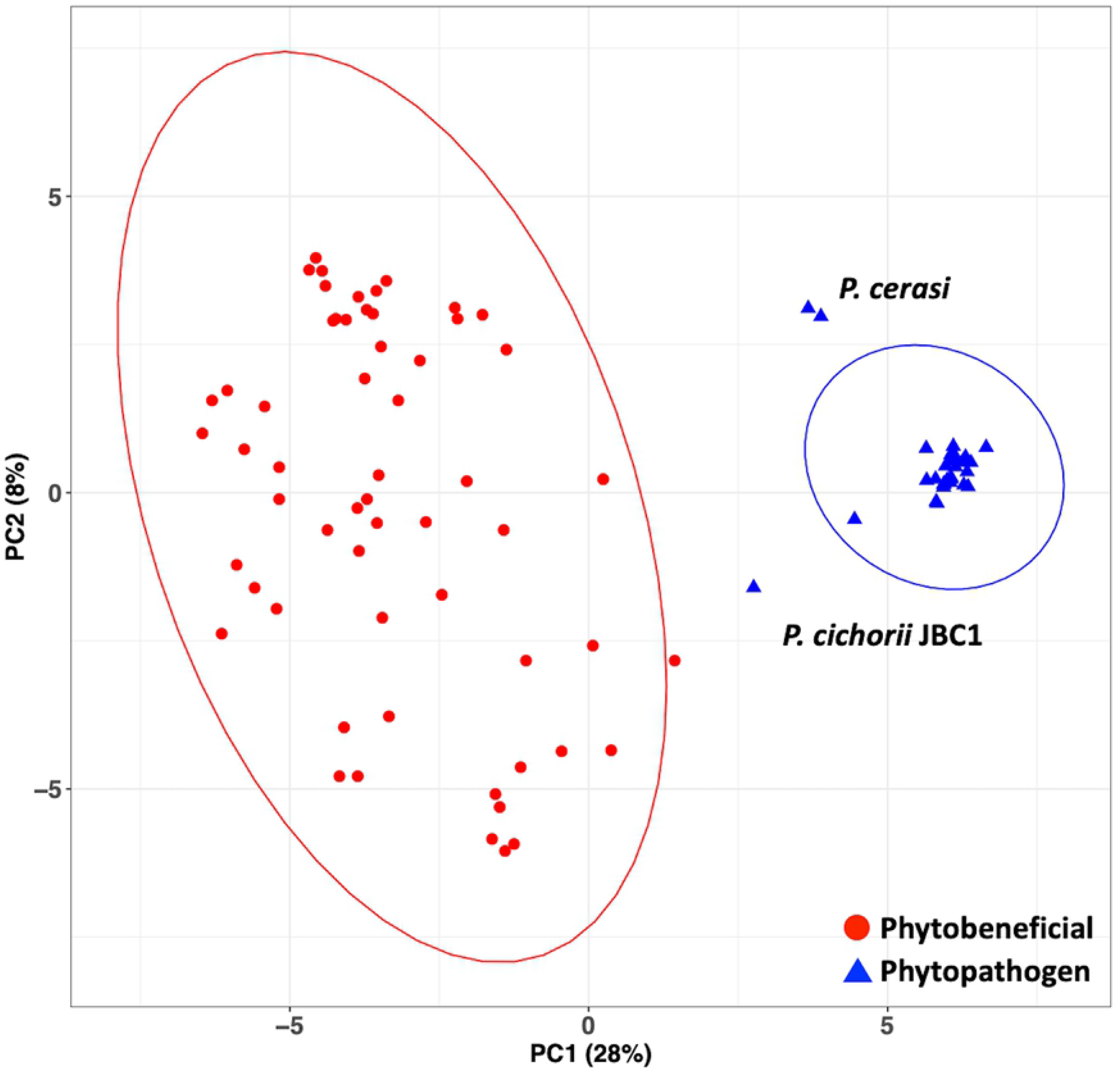
PCA based on GP-SND content as variables. The fraction of the variance is given in parentheses. *P. cichorii* JBC1 and two strains of *P. cerasi* are outside 95% confidence ellipse of the phytopathogenic group.

To further understand the contribution of each GP to the separation, we performed an enrichment analysis on the results obtained with the GP-PA, GP-SD and GP-SND approach (**S3 Table**). The enrichment analysis was performed on the binary data of presence and absence of the properties by considering “PARTIAL” as presence or absence separately, creating two enriched sets per approach. Subsequently, the two enriched sets were intersected to create the enriched set for that particular approach. Lastly, an overall enriched set was constructed by considering only the GPs that were enriched in the GP-SD and GP-SND approaches (**Table 2**).

**Table 2:** Genome Properties related to the plant-associated phenotype: enrichment analysis.

| Genome Property | Description | Adjusted P-value |
| --- | --- | --- |
| *GPs enriched in phytobeneficial strains* |  |  |
| GenProp0238* | 2-aminoethylphosphonate catabolism to acetaldehyde | $< 10^{-6}$ |
| GenProp0721* | 2-aminoethylphosphonate (AEP) ABC transporter, type II | $< 10^{-6}$ |
| GenProp0613* | Cytochrome c reductase | $< 10^{-6}$ |
| GenProp0907 | Poly-beta-1,6 N-acetyl-D-glucosamine system, PgaABCD type | $< 10^{-6}$ |
| GenProp0271 | Trehalose utilization | $< 10^{-6}$ |
| GenProp1745 | GA12 biosynthesis | $< 10^{-6}$ |
| GenProp1189 | MqsRA toxin-antitoxin complex | $< 10^{-6}$ |
| GenProp1645 | Zeaxanthin biosynthesis | $< 10^{-6}$ |
| GenProp0659 | Tryptophan degradation to anthranilate | $7.96 \times 10^{-5}$ |
| GenProp0895 | Alcohol ABC transporter, PedABC-type | $7.01 \times 10^{-4}$ |
| GenProp0902 | Quinohemoprotein amine dehydrogenase | $1.40 \times 10^{-3}$ |
| GenProp1516 | Phosphatidylcholine biosynthesis V | $5.37 \times 10^{-3}$ |
| *GPs enriched in phytopathogenic strains* |  |  |
| GenProp0908* | 2,3-diaminopropionic acid biosynthesis | $< 10^{-6}$ |
| GenProp0813* | Pyrimidine utilization | $< 10^{-6}$ |
| GenProp1165* | PhnGHIJKL complex | $< 10^{-6}$ |
| GenProp1381 | Methylphosphonate degradation I | $< 10^{-6}$ |
| GenProp0236 | Phosphonates ABC transport | $2.62 \times 10^{-3}$ |
| GenProp0710 | Generic phosphonates utilization | $2.62 \times 10^{-3}$ |
| GenProp1193 | RelBE toxin-antitoxin complex | $3.19 \times 10^{-2}$ |
| GenProp1566 | D-galactonate degradation | $3.64 \times 10^{-2}$ |

\*These Genome Properties are also important random forest features (Table 3).

To extend our analysis utilizing the full information of the classes and to capture feature importance, a Random Forest (RF) classifier was built. For 99% of the strains, the RF classifier correctly predicted the phenotype. The only exception was *Pseudomonas cichorii* JBC1, which had been reported to be pathogenic but was classified by RF-classifier as phytobeneficial. To study the discriminating variables further, variable selection from RF was implemented (**Table 3** and **S3 Table**). These variables were integrated with the list of enriched GPs to generate a comprehensive list of key genomic features contributing to the plant-associated phenotype (**Fig 4**).

**Table 3:** Genome Properties related to the plant-associated phenotype: Random Forest features importance.

| Genome Property | Description | Predictive power** |
| --- | --- | --- |
| GenProp0813* | Pyrimidine utilization | 500 |
| GenProp0908* | 2,3-diaminopropionic acid biosynthesis | 500 |
| GenProp0721* | 2-aminoethylphosphonate (AEP) ABC transporter, type II | 329 |
| GenProp0238* | 2-aminoethylphosphonate catabolism to acetaldehyde | 328 |
| GenProp0615 | Cytochrome c based oxygen reduction and quinone re-oxidation | 251 |
| GenProp0613* | Cytochrome c reductase | 243 |
| GenProp1629 | Propanoyl-CoA degradation I | 215 |
| GenProp1572 | L-carnitine degradation I | 145 |
| GenProp1562 | Fatty acid salvage | 53 |
| GenProp1717 | Fatty acid beta-oxidation I<br>(GenProp1308, GenProp1510 and GenProp1544) | 53 |
| GenProp1165* | PhnGHIJKL complex | 2 |
| GenProp1251 | L-tyrosine biosynthesis I | 2 |
| GenProp1281 | Hydrogen sulfide biosynthesis I | 1 |
| GenProp1681 | L-cysteine degradation III | 1 |

\*GP also found in the enrichment analysis. \*\*Numbers were obtained using recursive feature
elimination (500 iterations)

**Fig 4:**
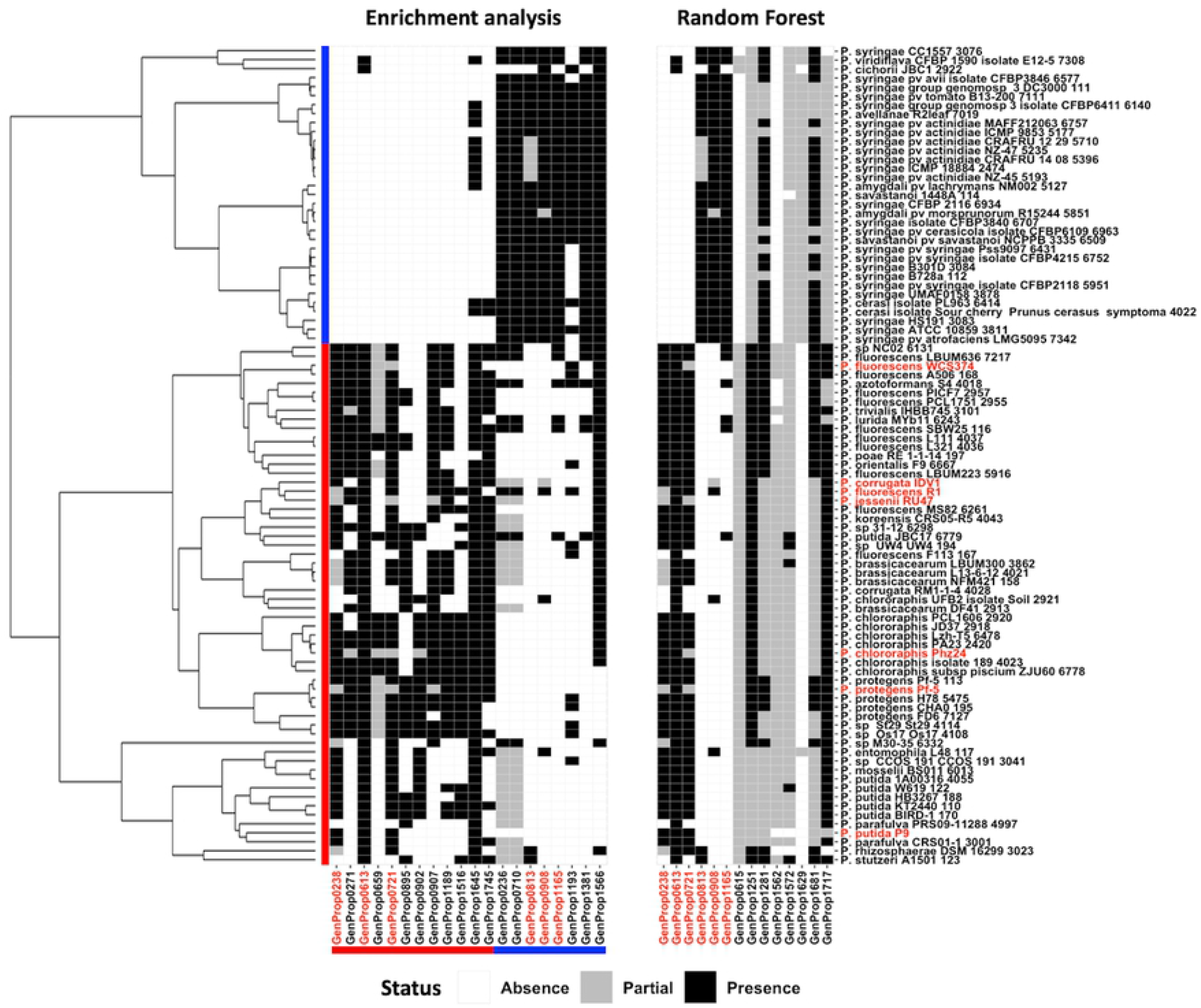
Representative list of discriminating Genome Properties obtained with the GP-SND approach. Left panel: enrichment analysis, right panel: Random Forest feature selection. Red lines indicate the phytobeneficial strains (vertical) and enriched traits (horizontal). Blue lines indicate the phytopathogenic strains (vertical) and enriched traits (horizontal). Newly sequenced strains are in red. Enriched GPs that were also highlighted in the RF feature importance analysis are indicated in red.

### Prediction validation

A set of 67 newly retrieved *Pseudomonas* genome sequences were analyzed for the presence of GPs using the GP-SND approach and used in RF performance evaluation (**S1 Table**). Confirming the capability of GP content to predict the plant-associated phenotype, a PCA of the full dataset (training and validation) indicated that the separation between the phytobeneficial and the phytopathogenic strains was retained. Additionally, a clustering of bioremediation strains with phytobeneficial strains was observed (**Fig 5Error! Reference source not found**.). Unclassified strain *Pseudomonas* sp. KBS0707 was positioned within the pathogen group. As all *P. syringae* are considered to be phytopathogenic, the unclassified *P. syringae* isolate inb918 was of interest as it appeared to be a phytobeneficial strain. The ANI score however suggested that strain inb918 might have been taxonomically misclassified as among the *P. syringae* strains the pair-wise score between this strain and the others remained below 79% (**Fig 5**). Lastly, the RF classifier was applied to the validation set and yielded the same predictions as the PCA.

**Fig 5:**
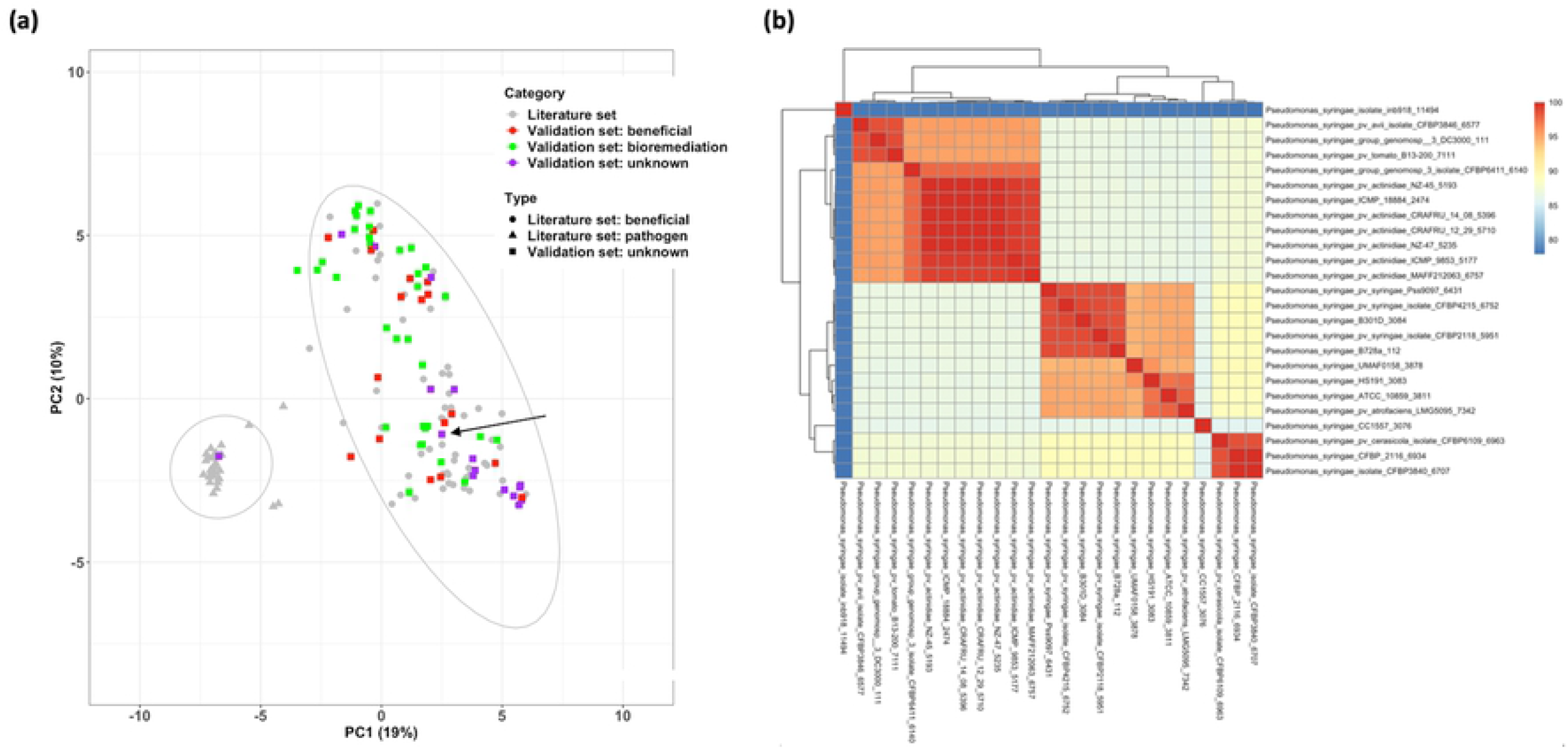
Analysis of the validation set. (a) PCA of the complete set of SND-GP data: variance is indicated in brackets. Previously analyzed *Pseudomonas* strains and previous obtained 95% confidence ellipses are in gray. The validation set is composed of 3 classes: phytobeneficial strains (red squares), bioremediation strains (green squares) and unclassified strains (purple squares). The arrow points at *P. syringae* isolate inb918. (b) Average Nucleotide Identity (ANI) score among *P. syringae* strains. *Pseudomonas syringae* isolate inb918 is at the top left.

## Discussion

Plants live in symbiotic interactions with microbial communities, which are complex networks composed of interacting microbiotic nodes. The sum of these interactions can be beneficial for plant growth and development, detrimental or neutral. Many important PGPR as well as plant pathogens belong to the genus *Pseudomonas*. The genomic diversity observed at species [22] and strain level suggests that Pseudomonas spp. have a broad potential for evolutionary adaptation to different environments. Consequently, the plant-associated lifestyle of a *Pseudomonas* strain is likely to be the result of a combinatorial accumulation and emergence of a diverse set of contributing traits. A selected isolated genome encoded feature therefore will have limited power to confidently predict the plant-associated phenotype.

Differences between phytopathogenic and phytobeneficial strains emerge at all levels of analysis. At genome sequence similarity level, a separation between the two groups was prominent. As most of the described phytopathogenic genomes in the scientific literature are *P. syringae* strains, a higher degree of sequence similarity was observed for the phytopathogenic group. The ANI score, however, does not take into account the most variable genomic regions that are likely to harbor genes that function in the biological relevant differences and would provide further insight in the functional diversity within the two phytotypes. The Genome properties (GP) annotation system was applied in this study to specifically address functional differences encoded in the genomes.

GPs represent not only metabolic pathways but also various other classes of functional attributes and provide, compared to KEGG and SEED, a better functional annotation coverage [21]. The GP annotation system is organized as a doubly linked rooted DAG. Leave nodes use domains as evidences, parent nodes, representing super-pathways, use leaf node GPs as evidences. For a functional genome comparisons at a larger scale, protein domains are better scalable and less sensitive to sequence variation compared to techniques based on sequence similarity [24]. By focusing on the reconstruction of domain-based GPs only, feature independence is promoted, and the complexity of the RF-model is reduced. In total 848 domain-based GPs were annotated to be (likely) present in one or more of the here studied *Pseudomonas* strain. Underpinning the genomic diversity of the 91 *Pseudomonas* strains used in this study, in contrast a functional core of maximal 154 complete and persistently present GPs was obtained. While for obvious reasons by far most of the typical eukaryotic GPs were not detected, a limited number of the *Pseudomonas* GPs may have some domain overlap with GPs of similar function typically found eukaryotic species. An example is the domain overlap between GenProp1717 and the “peroxisomal” GPs GenProp1308, GenProp1510 and GenProp1544 all involved in fatty acid beta-oxidation which we treated as one.

Three different approaches were used to determine the domain-based GP content of each strain. Implementation of the domain colocalization constraint mirrors the operonic structure common in bacterial genomes [25]. For domain colocalization a sliding window of 20 domains was chosen as it covers 1255 of the 1286 GPs (98%) with the most abundance group of GPs being GPs with two evidences (396 GPs) (**S5 Fig**). As the average domain copy number is 2.3, indicating that the same domain could be assigned to multiple functions across the genome, inclusion of protein domain colocalization in GP reconstruction also increases the prediction certainty of those GPs and further promotes the selection accessory traits, some of which may be acquired by lateral transfer, as RF-variables. Very similar results were obtained with GP-SND and the strain specific GP-SD method, suggesting that domain clustering most likely yields operonic structures.

The validation data was used to explore the performance of the RF classifier. For most validation data the RF firmly supports the discrimination between the beneficial and the pathogenic strains. *P. cichorii* JBC1 was classified as non-pathogenic. However, that does not directly translate into it being beneficial. **Fig 4** shows that *P. cichorii* JBC1 still contains three GPs associated with pathogenicity: ‘2,3-diaminopropionic acid biosynthesis’ (GenProp0908), ‘RelBE toxin-antitoxin complex’ (GenProp1193) and ‘D-galactonate degradation’ (GenProp1566). *P. cichorii* JBC1 has already been reported to be quite different to other pathogenic *Pseudomonas* at the genome level [26] and our results confirm this finding suggesting that there may be different mechanisms for pathogenicity associated with this strain.

RF recursive feature elimination and GP enrichment analysis was used to select a minimal set of GP-variables needed for a good prediction of the phenotype [27]. GenProp0238 and GenProp0721 are two of those important RF-variables and are shown to be enriched in phytobeneficial strains. The two GPs are related to mechanisms of phosphonate utilization, which have been shown to occur in *Pseudomonas* and also in other microorganisms [28]. Phosphonate is a form of phosphorus, which is essential for many biological processes [29]. However, both groups show differences in the usable form of phosphonate. Most phytobeneficial strains appear to be able to utilize only 2-aminoethylphosphonate (AEP) via the genome properties: ‘2-aminoethylphosphonate catabolism to acetaldehyde’ (GenProp0238) and ‘2-aminoethylphosphonate (AEP) ABC transporter, type II’ (GenProp0721), whereas the phytopathogens are able to access broader forms of phosphonates, as also shown by the enriched protein domain, via ‘phosphonates ABC transport’ (GenProp0236), ‘generic phosphonates utilization’ (GenProp0710), ‘PhnGHIJKL complex’ (GenProp1165) and ‘methylphosphonate degradation I’ (GenProp1381) [30]. AEP is the most abundant C-P compound in nature while other phosphonates and their derivatives are substances used in agriculture (herbicides, fungicides and insecticides) and pharmacy (antibiotics) [31]. It has been reported that the virulence of pathogenic species was enhanced under conditions of orthophosphate limitation [32]. Thus, we hypothesize this could be due to the presence of genome traits that enable them to access a wider set of phosphate sources.

GenProp0908 is another important RF-variable. This GP was found to be enriched in phytopathogenic strains and is involved in 2,3-diaminopropionic acid biosynthesis (DAP). DAP is a precursor of several secondary metabolites, such as siderophores, neurotoxins and antibiotics [33]. Pyoverdins, the principal siderophores, have been reported to be produced exclusively by the pathogens, such as *P. syringae* and *P. cichorii* [34]. Siderophores are important metabolites involved in iron acquisition [35]. Iron is crucial to many metabolic processes and is therefore required to maintain cells in a healthy state [36]. The stronger ability to scavenge for iron, and the phosphonate previously mentioned, will increase the fitness of the pathogens.

Two GPs strongly enriched among the phytobeneficial strains are GenProp0907, and GenProp0902. GenProp0907 represents a cluster of four genes involved in the synthesis, modification and export of the biofilm adhesin poly-beta-1,6-N-acetyl-D-glucosamine and the four domain evidences represent the four genes required. The GP is not present in the phytopathogen group and found to be complete as likely operonic structures in 39 phytobeneficial strains. Biofilms of the PgaABCD type have been studied in *Escherichia coli* [37] but not in *Pseudomonas* species. GenProp0902 represents quinohemoprotein amine dehydrogenase (QHNDH). QHNDH is a three-subunit enzyme located in the periplasmic space of *P. putida* and part of the amine oxidation respiratory chain. QHNDH catalyzes the oxidative deamination of primary amines when used as a sole carbon and energy source [38]. The GP consists of four evidences, three domains representing the alpha-, beta-and gamma-subunit of the enzyme and one representing the QHNDH maturation protein. This likely operonic GP was found to be complete in 24 biocontrol strains and not present in the pathogen group. As these GPs are only present in subset of the phytobeneficial strains, they did not emerge as important RF-variables in recursive feature elimination.

Protein domains associated with Type II secretion system (T2SS) were found to be enriched among the phytobeneficial strains while domains involved in the type III secretion system (T3SS) were found to be enriched among the phytopathogenic strains. T2SS is described by GenProp0053 and consists of 10 non-optional evidences and 3 optional domains. GP results however, indicated for both phytobeneficial and phytopathogenic strains a “PARTIAL” status for this GP. Similarly, the type III secretion system, represented by GenProp0052 is considered to be a key virulence factor and has been considered as evidence for pathogenicity in many genome studies [17,39,40]. GenProp0052 is a complex GP consisting of 14 evidences and 28 optional domains. Due to the set zero threshold for “PARTIAL” for this specific GP, a single evidence domain will already result in a “PARTIAL” status.

Eighteen protein domains enriched in phytopathogens are described to be involved in Type III secretion systems. Eleven of those enriched domains are used as evidences for GenProp0052. One other, TIGR02551, did also occur in the pathogen set but was considered not to be enriched after the Bonferroni adjustment. In contrast, the two missing evidences, TIGR02105 and TIGR02546 are only present in five phytobeneficial genomes. Thus, amongst the 91 Pseudomonas strains all 14 evidences are present, but none of the strains used in this study have the complete set of 14 evidences.

Due to the ‘Partial’ status of GenProp0052 (T2SS) and GenProp0053 (T3SS) for both phytotypes these GPs were not enriched, nor were they selected as discriminating variables in RF classification. We further examined the distribution of the GenProp0053 and of GenProp0052 evidences over all strains (**S6 Fig**). The distribution showed that protein domains linked to GenProp0052 more consistently occurred in the pathogen group with more variation in the phytobeneficial group. The result suggests that the abundance of T3SS related domain content could be sufficient for an indication of the pathogenicity. However, there is no guarantee that the feature is functional due to the missing evidences.

Specifically, for the phytobeneficial group a number of enriched GPs suggested a role for pathways involved in the degradation and utilization of trehalose (GenProp0271), tryptophan (GenProp0659) (Table 2), tyrosine (GenProp1251) and carnitine (GenProp1572) (Table 3). On the other hand, phytopathogenic strains appears to be more specialized in the degradation of galactonate (GenProp1566) and cysteine (GenProp1681). Carbon sources that were predicted to be degradable by preferably the phytobeneficial group could contribute to the agricultural industry. These substrates could be used as fertilizers, growth promotors, or as additives to alternate the microbial composition [41]. Similar to elicitors, which directly enhance plant defense and resistance, this indirect approach could be applied to the existing microbial community to select for the beneficial strains and potentially increase the productivity of the crop. [42]. On the other hand, carbon sources that might prolong saprobic growth and survival of pathogens should be avoided.

Other GPs found in the phytobeneficial group are linked to four ‘human hormones’, which are ‘mineralocorticoid biosynthesis’ (GenProp1644), ‘estradiol biosynthesis II’ (GenProp1417), ‘glucocorticoid biosynthesis’ (GenProp1666) and ‘pregnenolone biosynthesis’ (GenProp1740). The evidence shared by these hormones, domain PF00067 (cytochrome P450), is the same as for ‘GA12 biosynthesis’ (GenProp1745). Hence, only GA will be further discussed. Gibberellin 12 (GA_12_), is the common precursor of all gibberellins (GA) [43]. GA phytohormones play important roles in influencing the growth and development of the host plants [44] and GA from *Pseudomonas* could increase seed germination [45].

Not all known traits are represented by a GP. Many of those are found in phytopathogenic strains such as, coronatine, cytokinin and auxin [46]. We examined the presence of the protein domains associated to these traits in our dataset (**S7 Fig**). The results showed that the associated protein domains are generally present in both groups. Among these domains, only PF08659 and PF16197 were enriched in the phytopathogenic group. This suggests that the occurrence of these, known to be, phytopathogenic traits may not be sufficient as a genetic marker to identify the pathogenicity of a strain.

In conclusion, domain-based Genome Properties appear to be robust computational features to differentiate between phytobeneficial and phytopathogenic *Pseudomonas* strains and our analysis shows that incorporation of domain colocation further increases their relevance. By combining traditional statistical analysis (enrichment analysis) and machine learning methods (random forest) we were able to identify new discriminating genome properties that can be used to identify species that promote plant growth. These could be applied in strategies to develop synthetic PGPR communities and to formulate soil additives to improve plant health and performance.

## Materials and Methods

### Genome retrieval and annotation

*Pseudomonas* genomes with were downloaded from Pseudomonas Genome DB version 17.2. The validation set was obtained from database version 20.2 (https://www.pseudomonas.com) [22]. Genomes were manually categorized according their phytotype using literature data. Additionally, 7 genome sequences were (re)sequenced from phytobeneficial strains *P. putida* P9 (accession ERS6670306), *P. Corrugata* IDV1 (accession ERS6652532), *P. fluorescens* R1 (accession ERS6670181), *P. protegens* Pf-5 (accession ERS6652530), *P. chlororaphis* Phz24 (accession ERS6670416), *P. jessenii* RU47 (accession ERS6670307) and *P. fluorescens* WCS374 (accession ERS6652531). DNA was extracted using the Epicenter Masterpure kit (Epicentre Technologies, USA) according to the manufacturer’s protocol, quantified. For with the Infinite® 200 PRO (Tecan, Männedorf, Switzerland) using the Quant-iT™ PicoGreen™ dsDNA Assay Kit (ThermoFisher, Waltham, USA) according to the manufacturer’s protocol. The strains were sequenced on the PacBio Platform (Pacific BioSciences, Menlo Park, USA). A total of 4 ug DNA was sheared to 7 Kb and two SMRT bell libraries were prepared using the kit Barcoded Adapters for Multiplex SMRT sequencing in combination with the Sequel Binding Kit V2.0 and the Sequel Polymerase 2.0 Kit. Per library, a pool with sheared DNA of all strains was used as input according to the manufacturer’s protocol. Sequencing was done on a Sequel system operated at the services of Business Unit Bioscience, Wageningen Plant Research (Wageningen, The Netherlands). Subsequently, de-multiplexing was performed by aligning the barcodes to the sub-reads with pyPaSWAS version 3.0 [47]. Canu version 1.6 [48] was used to assemble the PacBio reads

The SAPP semantic annotation framework [49] was used to systematically (re)annotated the genomes. Briefly, protein encoding genes were de novo predicted using Prodigal 2.6.3 [50] and protein domains were characterized with InterProScan 5.36-75.0 using the Pfam and TIGRFAMs databases [51–53]. Annotation data and meta-data was stored in a semantic database using the GBOL ontology [54,55]. SPARQL queries were used to extract protein domain identifiers, and the location and direction of the corresponding gene.

### Data processing

OrthoANI version 1.40 was used to calculate the Average Nucleotide Identity (ANI) score for all genomes [56]. PygenProp, was used to infer from each genome domain-based GPs, [55]. Three criteria were applied; “PA”, considering only domain presence as evidence, “SND”, synteny-non-directional, requiring the genome location of the corresponding domains to be in close proximity and “SD” that in addition to gene location also considers strandness. For SND and SD a nearest neighbor approach and a sliding window of 20 protein domains was applied. Each GP was classified as either ‘YES’, or ‘PARTIAL’ according to the completeness of the set of evidences.

### Statistical analysis

The natural grouping of the data was visualized using principal component analysis (prcomp package). Then, with R packages; fisher.test and p.adjust, Fisher Exact Test with Bonferroni correction was applied to protein domains and the genome properties to test for enrichment. This analysis identified the over-and under-represented features. GP data was reassessed twice by considering ‘PARTIAL’ as either ‘YES’ or ‘NO’. The enriched list was created by intersecting the two cases of ‘PARTIAL’. Enrichments were considered significant if the adjusted p-value after Bonferroni correction of the GP is below 0.05.

The Random Forest classifier was created using R package randomForest v4.6-14 [58]. Labelled data were divided into training and test sets. The unbiased training set was created with equal numbers per group determined by using 75% of the smaller group, the pathogen group, resulting in 25 strains per group. Therefore, the test set remains with 33 phytobeneficials and 8 phytopathogens. The Variable Selection from Random Forests v 0.7-8 (varSelRF) package in R was used to determine variable importance. We used 5000 trees for the first forest and 2000 trees for all additional forests during the iteration. Vars.drop.frac, the portion of the variable that is excluded on each iteration, was set to 0.2.

## Acknowledgements

WP is financially supported by a Royal Thai Government Scholarship, Thailand. TL acknowledges the support by the Dutch Ministry of Economic Affairs in the Topsector Program “Horticulture and Starting Materials” under the theme “Plant Health” (project number: TU 16022) and its partners (NAK, Naktuinbouw and BKD). PS and MSD acknowledge the Dutch national funding agency NWO, and Wageningen University and Research for their financial contribution to the Unlock initiative (NWO: 184.035.007).

## Supporting information

**S1 Table. List of strains.**

List of strains used in this study. The dataset used for the initial analysis and the validation data are in different tabs. The list provides strain’s name, their classification along with their corresponding annotation information.

(XLSX)

**S2 Table. Enriched protein domains.**

Enriched protein domains on phytopathogenic and phytobeneficial strains with the p-value and number of occurrences.

(XLSX)

**S3 Table. Genome Properties analysis.**

Genome Properties analysis results divided into 9 sheets. First three sheets are according to the analysis approaches: GP-PA (presence-absence), GP-SD (synteny-directional) and GP-SND (synteny-nondirectional). Sheets 4 and 5 are the enriched GP of the phytopathogen and beneficial respectively. Sheets 6 to 8 are the variable selection using the Random Forest using 3 analysis approaches. The final sheet are the GPs that are not presented according to any approaches.

(XLSX)

**S4 Fig. PCA using 3 approaches**.

(PDF)

**S5 Fig. Distribution of number of evidences of the Genome Properties**.

(PDF)

**S6 Fig. Distribution of non-optional evidences of GenProp0053 and GenProp0052**.

(PDF)

**S7 Fig. Presence and absence of protein domains associated to genes related to selected pathogenic traits found in *P. syringae***.

(PDF)

